# Early-life exposures to specific commensal microbes prevent type 1 diabetes

**DOI:** 10.1101/2024.02.15.580579

**Authors:** Jamal Green, John Deschaine, Jean-Bernard Lubin, Julia N. Flores, Sarah Maddux, Tereza Duranova, Logan Grimes, Paul J. Planet, Laurence C. Eisenlohr, Michael A. Silverman

**Affiliations:** Division of Infectious Disease, Department of Pediatrics, The Children’s Hospital of Philadelphia, Philadelphia, PA 19104, USA; Perelman School of Medicine, University of Pennsylvania, Philadelphia, PA 19104, USA; AMGEN, South San Francisco, CA 94080, USA; Institute for Immunology and Immune Health, Perelman School of Medicine, University of Pennsylvania, PA 19104, USA; Department of Microbiology, Perelman School of Medicine, University of Pennsylvania, Philadelphia, PA 19104, USA

**Author notes:** **Corresponding author:** Michael A. Silverman.

## Abstract

Early-life disruptions of the gut microbiome have long-lasting impacts on the risk of developing autoimmune diseases. How the composition of the early-life microbiota contributes to autoimmunity and whether manipulating it can prove therapeutically beneficial remains largely unexplored. Here we demonstrate that a simple consortium of nine early-life commensal bacteria (PedsCom) prevents type 1 diabetes (T1D) in diabetes-susceptible NOD mice. Remarkably, we find that this protection is completely dependent upon early-life colonization. During this critical time window of early-life colonization and immune development, specific microbes unexpectedly translocate from the gut to peripheral tissues and induce the tolerogenic responses required for T1D protection. These findings highlight how the timing and localization of microbial interactions during a pivotal stage of immune development contribute to protection from T1D. Altogether, these findings suggest an opportunity to develop microbial therapies for human infants to prevent autoimmune diseases.

**One sentence summary:** A defined consortium of early-life microbes shapes immune development and prevents type 1 diabetes.

## Introduction

Type 1 diabetes (T1D) is a T-cell-mediated autoimmune disease in which the insulin-producing beta cells in the pancreatic islets of Langerhans are destroyed. Unfortunately, the worldwide incidence of T1D continues to rise, with an especially rapid increase in children (*1–4*). While there is a strong genetic contribution to the disease, the rising incidence of T1D over the past few decades provides compelling evidence that environmental factors, such as the microbiome, also impact the risk of developing T1D (*5–7*).

The timing of microbial interactions with the developing immune system profoundly influences long-term immune health (*8–10*). In humans, the microbiome dramatically shifts in composition, diversity, and function during the first few years of life, after which it matures into an adult microbiome (*11*). In mice, this developmental process is more rapid, with mice initiating solid food around days 12-14, weaning from milk by day 21, and developing a microbiome that resembles that of an adult by the end of the 4^th^ week of life (*12–14*). These dramatic changes in the microbiome during weaning correspond with the remarkable development of components of the immune system that are essential for maintaining long-term immune homeostasis, including the development of peripheral regulatory T cells (pT_regs_) (*15, 16*), systemic anti-commensal IgG antibodies (*17, 18*), unconventional T cells (*19, 20*), a transition from maternal IgA to endogenously produced IgA (*21, 22*), and an induction of antigen-specific tolerance to commensal microbes and food antigens (*23, 24*). In addition, during the weaning period key components of gut barrier function, such as the level of mucosal IgA (*21*), the mucin gene *Muc2* (*25*), goblet-cell-associated antigen passages (GAPs) (*23*), and expression of toll-like receptors (TLRs) (*26*) are dynamically regulated. The importance of the early-life interactions between commensal microbes and the immune system is exemplified by the “weaning reaction” (*14*), a complex set of time-restricted, commensal-driven immune responses that occur around weaning. The weaning reaction, which includes a spike in inflammatory cytokines, opening of colonic GAPs to allow antigen sampling of the microbiome, and the development of pT_regs_, is required for the development of long-term immune health. Indeed, disruption of the microbiota, specifically around weaning, leads to profound susceptibility to immune dysfunction later in life (*14*). In humans, early-life perturbations of key host-microbe interactions are associated with an increased risk of diseases such as allergy, asthma, psoriasis, obesity, and inflammatory bowel disease (*9, 10, 27*). Epidemiologic and experimental studies suggest that the same is true for T1D (*28–31*). Altogether, the timing of specific microbial exposures is critical for educating a healthy immune system and may be critical for preventing T1D.

To rigorously study the interactions between early-life microbes and the developing immune system, we previously designed the Pediatric Community (PedsCom) which is a defined nine-member bacterial consortium that recapitulates the composition and functions of the early-life microbiome (*13*). This microbial community was derived from the intestinal microbiota of preweaning 14-day-old MHCII E-expressing NOD mice (Eα16/NOD). Eα16/NOD mice were selected as the source of microbiota because they effectively model human leukocyte antigen (HLA) class II haplotypes that protect humans from T1D (*32, 33*) and because early-life commensal microbiota from Eα16/NOD mice prevent the development of T1D in NOD mice (*29*). PedsCom covers >90% of the microbial reads across the small intestine, cecum, and large intestine and mirrors the phylogenetic diversity and functions of a complete pre-weaning microbiome. We previously used PedsCom to demonstrate that microbiome maturation supports immune system development in C57BL/6 mice (*13*).

In this study, we generated NOD mice colonized with PedsCom to experimentally test the impact of the early-life microbiome on the development of T1D. We found that the rationally-designed PedsCom consortium of early-life microbes induces IL-10-producing regulatory T cells in the intestine and pancreatic lymph nodes while restraining inflammatory CD4^+^ T-cell-mediated IFNγ production in the pancreatic islets of NOD mice. Most importantly, we identified a defined group of commensal microbes that prevent T1D by imprinting the immune system during a critical early-life window of development.

## Results

### Early-life microbial consortium prevents type 1 diabetes

Adult germfree NOD mice were colonized by oral gavage with a mixture of PedsCom microbes (10^9^ CFU of each member) (Table 1). After the initial colonization, NOD mice vertically transferred all nine of the PedsCom microbes to their progeny over multiple generations. As expected, the two *Lactobacillaceae* species predominated in the small intestine while *Parabacteroides distasonis* and the *Enterobacteriaceae, Kosakonia cowanii,* constituted >50% of the relative abundance in the cecum and large intestine (**Fig. 1A-B**). We also generated the Complex Mature Community (CMCom), a complex community derived from the cecal contents of a 6-week-old Eα16/NOD mouse, to model the adult microbiome (**fig. S1**). Germfree, PedsCom-, and CMCom-colonized NOD mice were bred and housed in separate gnotobiotic isolators to maintain their gnotobiotic communities.

**Figure 1.**
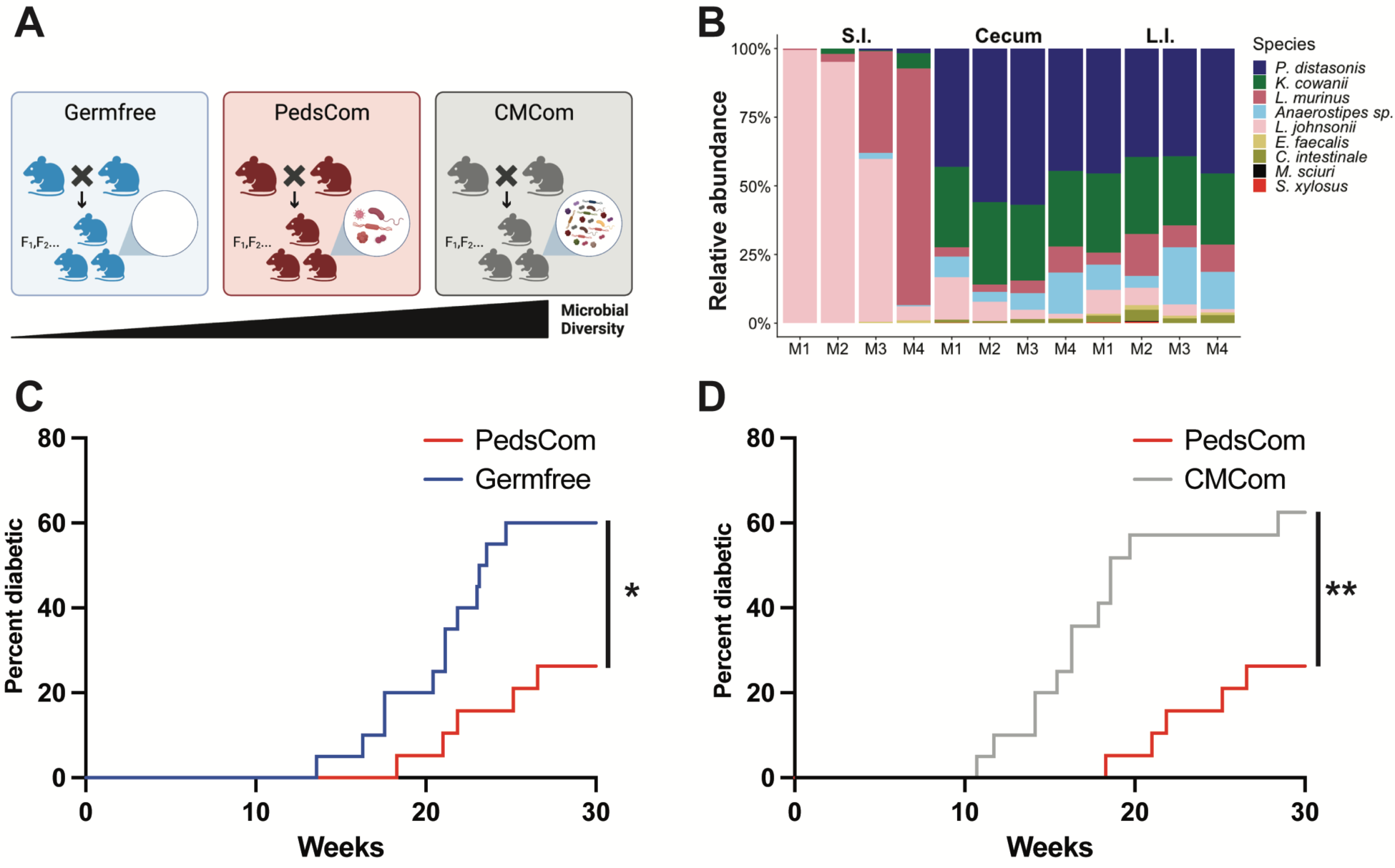
PedsCom colonization prevents diabetes in NOD mice. **A.** Schematic of gnotobiotic mice raised in individual gnotobiotic isolators: germfree, Pediatric Community (PedsCom), Complex Mature Community (CMCom). **B.** Relative abundance of PedsCom members in the small intestines, ceca, and large intestines of adult PedsCom-colonized NOD females. M1, M2, M3, and M4 denote individual mice. **C-D.** Diabetes incidence in NOD female mice born to PedsCom dams (n=19), germfree dams (n=20), or CMCom dams (n=19). Groups compared by the Log-rank test, * p< 0.05, ** p< 0.01.

**Table 1.**
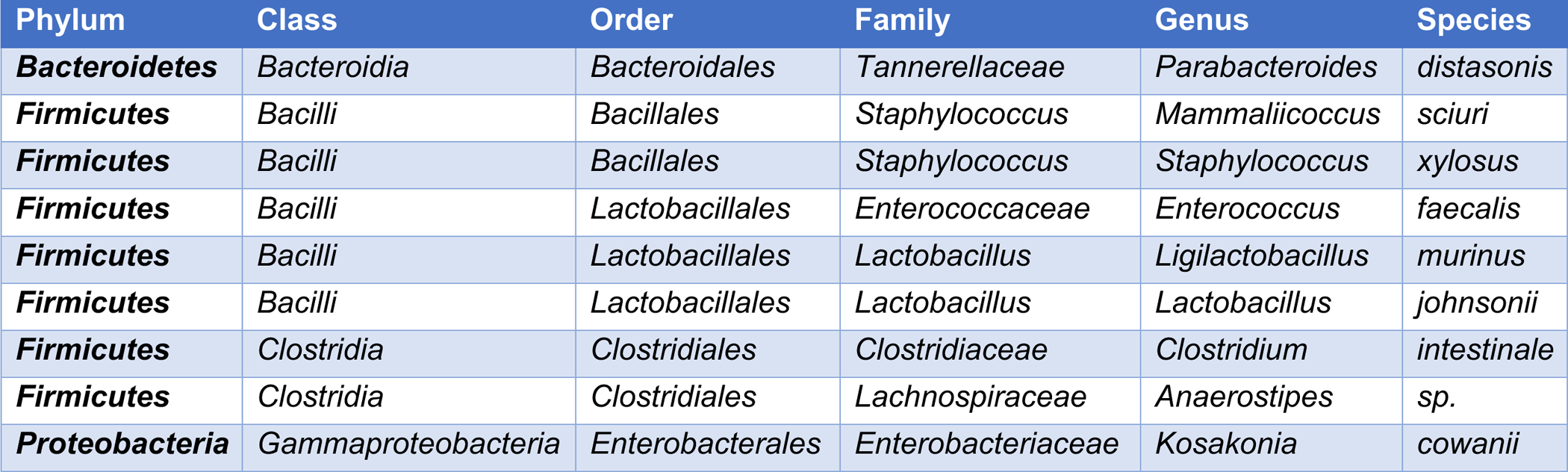
Pediatric Community (PedsCom)

To determine the extent to which early-life microbes derived from diabetes-protected Eα16/NOD mice impact the development of T1D in diabetes-susceptible mice, we compared the incidence of diabetes in PedsCom-colonized NOD mice to the incidence of diabetes in germfree NOD mice and in CMCom-colonized NOD mice. Since early-life exposure to commensal microbes shapes immune system development, we used PedsCom and CMCom NOD mice that received their microbiota by natural vertical transmission from their dam. The incidence of diabetes in female PedsCom NOD mice was significantly lower than in germfree NOD mice (**Fig. 1C**; 26.3% vs 60.0%, p<0.05), demonstrating that PedsCom microbes provide protection from developing diabetes. Remarkably, PedsCom microbes also significantly decreased the incidence and delayed the onset of diabetes compared to CMCom NOD mice (**Fig. 1D**; 26.3% vs 63.1%, p<0.01). Thus, colonization from birth with the nine early-life PedsCom microbes provides greater protection from diabetes than an entire complex adult-derived intestinal community. The lower diabetes incidence in PedsCom-colonized NOD mice compared to germfree and CMCom NOD mice demonstrates that this nine-member bacterial consortium is enriched for commensal microbes that are adept at preventing T1D.

### PedsCom colonization induces weaning-associated peripheral regulatory T cells and restrains IFNγ in the pancreatic islets

To determine the immunologic impacts of PedsCom colonization in NOD mice, we investigated CD4^+^ T cell populations in gut tissues and systemic sites relevant to T1D. Specific microbes and microbial communities can restrain inflammation through the induction of regulatory CD4^+^ T cells (*34, 35*). Microbe-induced pT_regs_ that develop during the 3^rd^ and 4^th^ weeks of life are hallmarks of weaning-associated immune development in the murine gut. They are identified by expression of transcription factors (Foxp3^+^, RORγ^+^, and Helios^−^) and are most abundant and well-studied in the intestinal lamina propria (*15, 16*). More recently, pT_regs_ have been reported to restrain insulitis (*36, 37*) and the development of T1D (*36*) in NOD mice.

PedsCom colonization induced a significantly higher proportion of pT_regs_ in the lamina propria of the cecum and large intestine, the mesenteric and pancreatic lymph nodes (MLNs and PLNs), and the spleen compared to germfree NOD mice (**Fig. 2A-B** and **fig S2A**), indicating that PedsCom stimulates the development of pT_regs_ in the gut and systemic sites. In contrast, when compared to CMCom-colonized mice, PedsCom microbes induced a lower proportion of pT_regs_ in the cecum and large intestine and a similar proportion of pT_regs_ in the PLNs and MLNs (**Fig. 2A-B and fig S2A**). Based on the development of pT_regs_ in the gut and systemic sites, we conclude that PedsCom colonization stimulates intestinal and systemic immune responses. Since PedsCom mice have comparable or lower levels of intestinal pT_regs_ than CMCom mice, the relative abundance of intestinal pT_regs_ does not adequately account for the different incidences of T1D in germfree, PedsCom, and CMCom NOD mice.

**Figure 2.**
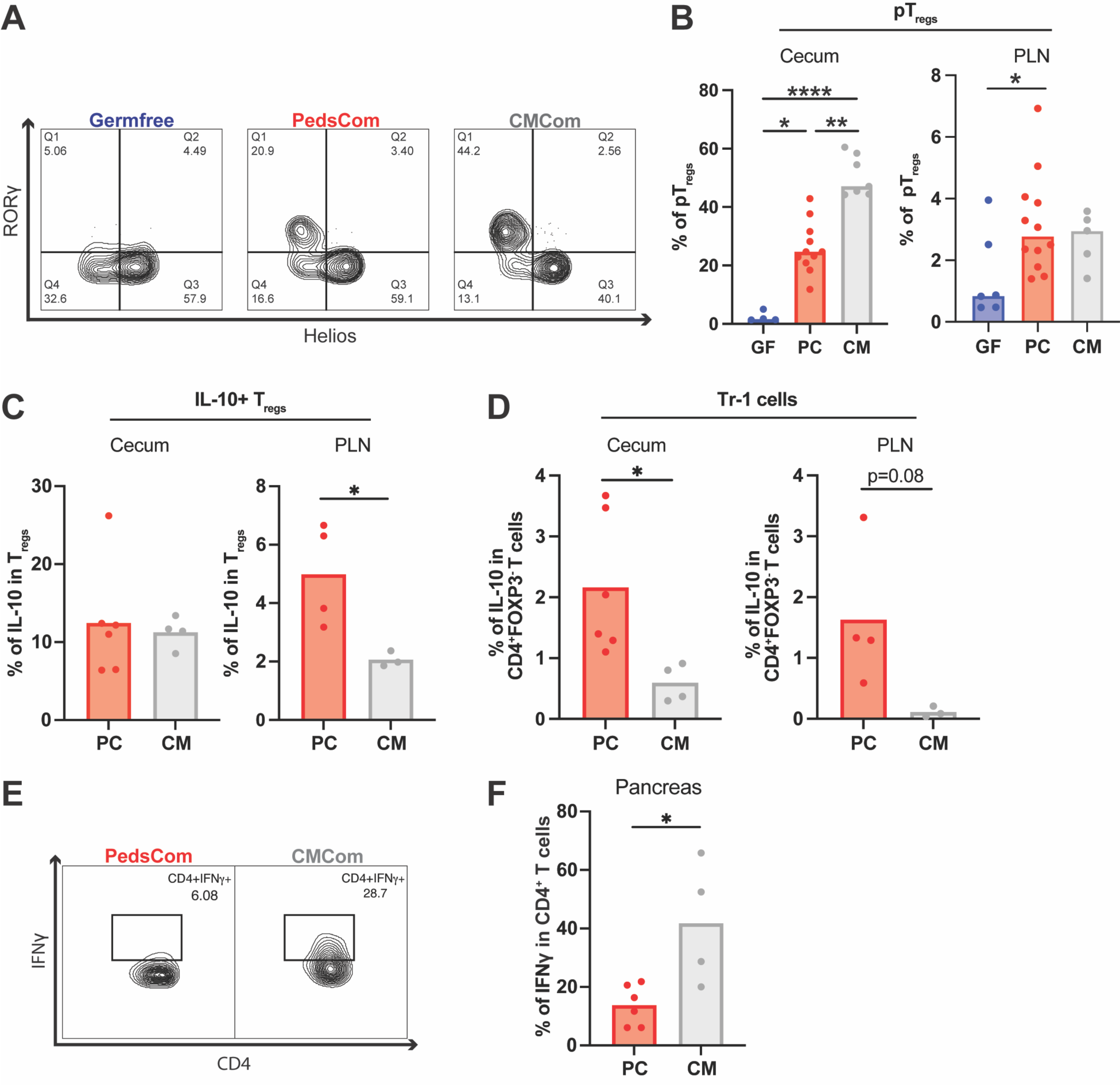
PedsCom induces pT_regs_ in the gut and restrains IFNγ in the pancreas. **A.** Representative flow cytometry plots of cecal CD4^+^FOXP3^+^RORγ^+^Helios^−^ pT_regs_. Cells were gated on CD45^+^TCRβ^+^CD4^+^FOXP3^+^cells. **B.** Percentages of pT_regs_ of total FOXP3^+^ T_regs_ in the ceca and PLNs of germfree, PedsCom, and CMCom mice. **C.** Percentages of IL-10-producing T_regs_ in the ceca and PLNs of PedsCom and CMCom mice. **D.** Percentages of IL-10 producing CD4^+^FOXP3^−^ T cells in the ceca and PLNs of PedsCom and CMCom mice. **E.** Representative flow cytometry of IFNγ-producing CD4^+^ T cells. **F.** Percent of IFNγ-producing CD4^+^ T cells in the pancreatic islets of PedsCom and CMCom mice. Data in panels A and B are representative of 6 experiments. Groups are compared with the Kruskal-Wallis test. n≥4 samples per tissue per gnotobiotic community, ages 5-11 weeks old. GF=germfree, PC=PedsCom, CM=CMCom. Data in panels C-F are representative of 4 experiments. Groups are compared with the Welch’s t-test, n≥3 tissues per gnotobiotic community, ages 10-12 weeks. *p<0.05, **p<0.01, ****p<0.0001.

We next compared IL-10 production, as a marker of T cell suppressive function, in the gut and the PLNs of germfree, PedsCom, and CMCom NOD mice. IL-10 restrains the development of T1D in some contexts via Foxp3-positive T_regs_ and Foxp3-negative type 1 regulatory (Tr1) cells (*38, 39*). Since specific commensal microbes robustly induce IL-10 production (*40–42*), we investigated the extent to which PedsCom colonization impacts IL-10 secretion from these regulatory cell populations in gut-associated and systemic sites relevant to T1D. Immune cells from germfree, PedsCom, and CMCom NOD mice were stimulated with phorbol myristate acetate (PMA)/ionomycin and then intracellularly stained for IL-10. PedsCom microbes induced a higher proportion of intestinal IL-10-producing CD4^+^ T cells and T_regs_ compared to GF mice and a similar proportion of intestinal IL-10-producing CD4^+^ T_regs_ compared to CMCom mice (**Fig. 2C and fig. S2B-C**). We conclude that microbe-induced pT_regs_ produce the majority of intestinal IL-10 since most of the IL-10-producing T_regs_ expressed the transcription factor RORγ (**fig. S2B-C**). These findings demonstrate that PedsCom microbes induce Foxp3-positive CD4^+^ T cells that produce the regulatory cytokine IL-10 in the gut and PLNs. PedsCom mice had a higher proportion of IL-10-producing T_regs_ in the PLNs compared to CMCom mice (**Fig. 2C**). This is notable as the PLN is a critical regulatory site for the initiation and progression of islet autoimmunity (*43, 44*).

Since PedsCom microbes robustly induced IL-10-producing Foxp3-positive CD4^+^ T cells, we next explored the extent to which this microbial consortium induces Tr1 cells, Foxp3-negative, IL-10-producing CD4^+^ T cells that can also be induced by specific commensal microbes (*40*) and contribute to protection from T1D (*38, 39*). Intestinal Tr1 cells migrate to the pancreas, decrease CD4^+^ IFNγ, and delay the onset of T1D in NOD mice through the actions of the IL-10 receptor (*38*). PedsCom mice had a higher proportion of Tr1 cells in the gut and PLNs compared to CMCom mice (**Fig. 2D**). These results demonstrate that PedsCom microbes promote the development of Tr1 cells as well as IL-10-producing Foxp3-positive T_regs_ which are potential immune pathways by which PedsCom microbes elicit protection from T1D.

IL-10 counterbalances Th1-driven autoimmunity (*45, 46*), in which IFNγ from CD4^+^ T cells sensitizes beta cells for CD8^+^ T-cell-mediated death (*47, 48*). Having established that PedsCom microbes induce IL-10-producing-regulatory-T cells in the cecum and PLNs at equal or higher proportions than CMCom, we hypothesized that PedsCom would decrease the proportion of pro-inflammatory IFNγ-producing CD4^+^ T cells in the pancreatic islets. Indeed, PedsCom mice had a dramatically lower proportion of IFNγ-producing CD4^+^ T cells in the pancreatic islets compared to CMCom mice (**Fig. 2E-F**, 14.1% vs. 40.6%, p<0.05). This restraint of IFNγ-producing CD4^+^ T cells in PedsCom NOD mice is evident when inflammation in the islets is fully developed but before the onset of clinical signs of T1D. However, PedsCom and CMCom mice have similar levels of insulitis at 10 weeks of age, despite striking differences in the risk for progression to T1D (**fig. S3**). As such, the increased proportion of IFNγ-producing CD4^+^ T cells in CMCom islets is consistent with more pathogenic islet autoimmunity. Together, these findings provide evidence that PedsCom microbes induce regulatory cell populations, including higher proportions of IL-10-producing-T_regs_ and -Tr1 cells compared to CMCom, restrict Th1 inflammation in the pancreatic islets, and prevent progression from insulitis to diabetes.

### *Parabacteroides distasonis*, *Anaerostipes* sp., and *Clostridium intestinale* induce regulatory T cells and are required to prevent T1D

One of the strengths of the PedsCom gnotobiotic model is the ability to deconstruct the community and determine the impact of each member on the development and function of the immune system. We hypothesized that specific PedsCom microbes induce regulatory cells that restrain autoimmunity. To test this hypothesis, 3-week-old germfree mice were orally gavaged with 10^8^ CFUs of cultured individual PedsCom members (**Fig. 3A**). After two weeks, we assessed the proportion of pT_regs_ in colonized mice. Monocolonization with specific microbes (*P. distasonis* and *Anaerostipes* sp.) induced pT_regs_ in the large intestine, cecum, and MLNs in NOD mice. In addition, *C. intestinale* and *K. cowanii* induced a modest but significant increase in the proportion of pT_regs_ in the cecum (**Fig. 3B-C**). The other five PedsCom microbes did not induce pT_regs_ in monocolonized mice. These experiments provide compelling evidence that *P. distasonis* and *Anaerostipes* sp. are the strongest *in vivo* inducers of intestinal pT_regs_ in PedsCom-colonized NOD mice.

**Figure 3.**
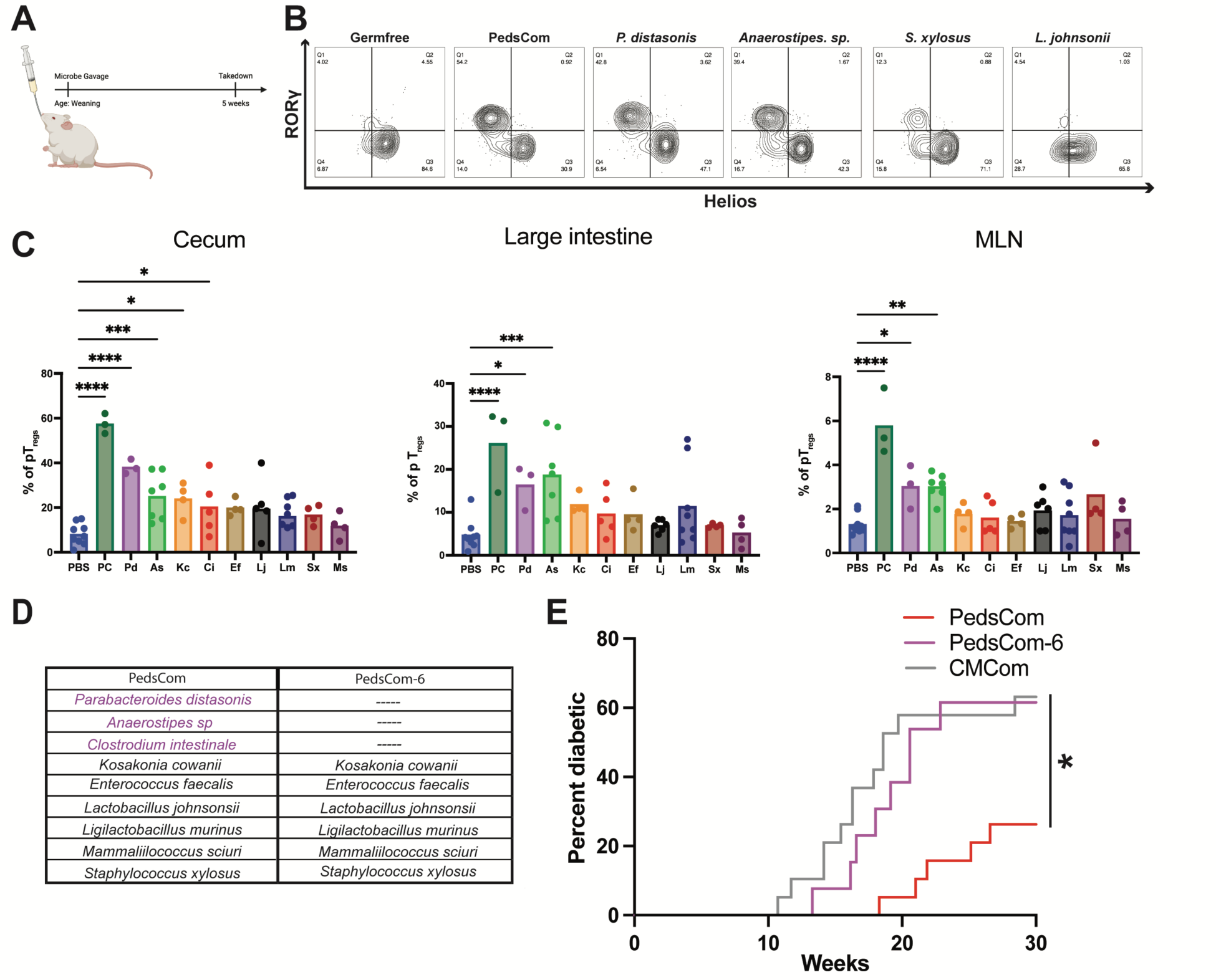
Specific PedsCom members induce pT_regs_ and prevent T1D. **A.** Schematic of PedsCom monocolonization experiments. Three-week-old germfree NOD mice were colonized with individual PedsCom species or with all nine PedsCom (PC) species and analyzed two weeks later. **B.** Representative flow cytometry plots of cecal CD4^+^FOXP3^+^RORγ^+^Helios^−^ pT_regs_ from germfree and monocolonized NOD mice. **C.** Percentages of pT_regs_ of total FOXP3^+^ T_regs_ in the ceca, large intestines, and mesenteric lymph nodes (MLNs) of monocolonized NOD mice. Groups were compared by ANOVA. n ≥ 3 mice per PedsCom microbe. **D.** Comparison of PedsCom and PedsCom-6 community (PedsCom without *P. distasonis, C. intestinale,* and *Anaerostipes sp*.). **E.** Diabetes incidence in NOD female mice born to PedsCom dams (n=19), CMCom dams (n=19), or PedsCom-6 dams (n=12). Groups were compared by the Log-rank test. *p<0.05, **p<0.01, ***p<0.001, and ****p<0.0001.

To further interrogate how PedsCom colonization impacts the immune system, we investigated the systemic antibody responses against each PedsCom member. Since T cells are required to develop systemic IgG1 antibodies against commensal microbes (*49*), we hypothesized that binding of the IgG1 subclass to specific PedsCom species would identify those microbes capable of inducing other T-cell responses, such as induction of pT_regs_. Since commensal microbes also induce T-cell-independent systemic IgG subclasses (IgG2b and IgG3) (*18*), we investigated whether PedsCom species induced these systemic antibodies as well. We performed microbial flow cytometry (mFLOW) to determine the degree to which IgG subclasses bind to pure cultures of each PedsCom member (*18, 49, 50*). To control for nonspecific binding, we also tested sera from germfree and RAG2 deficient mice (**fig. S4A-D**). PedsCom colonization induced IgG1 antibodies that bound to *Lactobacillus johnsonii, Staphylococcus xylosus,* and *Mammaliicoccus sciuri* (formerly *Staphylococcus sciuri)* (**fig. S4B)***. Enterococcus faecalis* demonstrated inconsistent IgG1 binding. *Mammaliicoccus sciuri* and *K. cowanii* elicited anti-commensal IgG2b antibodies (**fig. S4C)**. Anti-commensal IgG3 antibodies were not detected in PedsCom mice (**fig. S4D**). In contrast, we did not detect any systemic antibodies against *P. distasonis, Anaerostipes* sp., or *C. intestinale*. Of note, we were unable to assess *Ligilactobacillus murinus* by this assay due to high nonspecific binding. In summary, PedsCom-colonized NOD mice generate robust anti-commensal IgG1 and IgG2b antibodies against *Lactobacillus johnsonii, Staphylococcus xylosus, K. cowanii,* and *Mammaliicoccus sciuri*. In contrast to our initial hypothesis, the PedsCom microbes that induced pT_regs_ (*P. distasonis, Anaerostipes* sp. and *C. intestinale*) did not induce any systemic anti-commensal antibodies (**fig. S5A-B**). These data support a model in which a subset of microbes induce systemic IgG antibodies and a non-overlapping subset of PedsCom microbes induce pT_regs_.

Another powerful feature of the PedsCom gnotobiotic model is the ability to determine the impact of specific microbes by reconstructing the community without those microbes. Since *P. distasonis, Anaerostipes* sp., and *C. intestinale* induced pT_regs_, did not induce systemic antibodies, and are epidemiologically associated with T1D (*6, 51–54*), we hypothesized that one or more of these three microbes is necessary for the PedsCom consortium to protect from diabetes. To directly test this hypothesis, we generated a new six-member community, PedsCom-6, that lacks *P. distasonis, Anaerostipes* sp., and *C. intestinale* (**Fig. 3D** and **fig. S6**). As with the establishment of PedsCom, adult germfree NOD mice were gavaged with PedsCom-6 microbes, and their progeny assayed for the development of diabetes. We then compared the incidence of T1D between PedsCom-6, PedsCom, and CMCom NOD mice. PedsCom-6 completely lost the protective qualities of the nine-member PedsCom consortium (**Fig. 3E**). This loss-of-function gnotobiotic experiment demonstrates that *P. distasonis, Anaerostipes* sp., and/or *C. intestinale* are required for the protection from T1D conferred by PedsCom microbes.

### PedsCom microbes prevent T1D during an early-life window of development

The timing of microbial exposures during immune ontogeny regulates critical components of the immune system that have long-term impacts on the risk of developing inflammatory and allergic diseases (*14, 24, 55*). As such, we directly tested whether early-life colonization with PedsCom microbes is required for microbial protection from T1D. Germfree 6-week-old female NOD mice were cohoused with PedsCom mice to generate adult PedsCom NOD mice that had not been colonized with PedsCom microbes during early life (**Fig. 4A**). Remarkably, these NOD mice colonized by PedsCom after weaning developed T1D at a significantly higher rate than NOD mice colonized by vertical transfer from their dams (75.0% vs 26.3%) (**Fig. 4B**). The incidence of T1D in adult-colonized mice is similar to that of NOD mice colonized with CMCom microbiota at birth (63.1%) or maintained germfree (60.0%) (**Fig. 1C**). Importantly, the loss of protection in adult-colonized PedsCom mice was not due to differences in microbial colonization, as adult-colonized PedsCom NOD mice possessed similar proportions of microbes compared to mice born to PedsCom-colonized dams (**Fig. 4C**). These findings directly demonstrate that PedsCom microbes exert their protective effect during early life.

**Figure 4.**
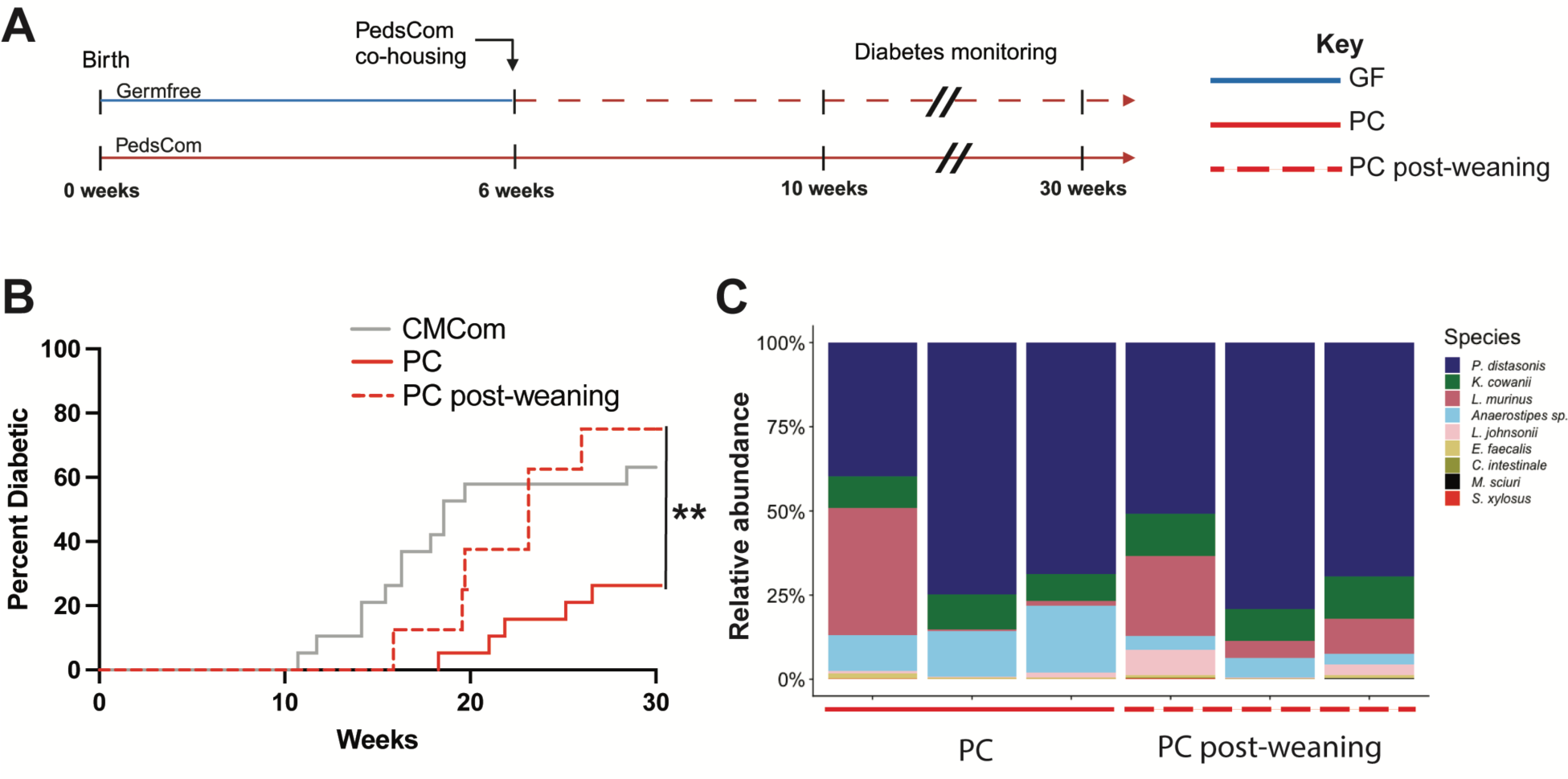
Protection from diabetes requires early life colonization with PedsCom. **A.** Schematic of PedsCom post-weaning colonization. **B.** Diabetes incidence in NOD female mice born to PedsCom colonized dams (n=19), germfree female mice colonized with PedsCom at 6 weeks of age (n=8), or NOD female mice born to CMCom colonized dams (n=19). Groups were compared by the Log-rank test. **C.** Composition of PedsCom members in fecal samples from PedsCom NOD mice colonized at birth and germfree NOD mice colonized with PedsCom at 6 weeks of age. GF=germfree, PC=PedsCom. **p < 0.01.

### Weaning-associated microbial translocation induces distinct immune responses

Since PedsCom microbial protection from T1D requires early-life exposure, we investigated whether PedsCom microbes potentially drive protection by gaining access to extra-intestinal sites during the tolerogenic window around weaning. We hypothesized that commensal bacteria translocate from the gut to other sites at weaning since young mice are still developing components of the intestinal barrier such as endogenous IgA (*21*), M cells (*25*), and the intestinal mucus layer (*56*). While a “leaky gut” contributes to the development of T1D in some contexts (*57*), there is limited direct evidence that a dysfunctional intestinal barrier early in life initiates this autoimmune disease. In contrast, several studies demonstrate that early-life commensal microbes induce immune responses, such as pT_regs_ (*23, 58*) and mucosal IgA (*22*), which are important for developing tolerance and preventing autoimmunity. We aseptically collected and cultured tissue homogenates from the MLNs, liver, and spleen, which represent routes of lymphatic, portal vein, and systemic translocation, respectively (*59, 60*). Live PedsCom microbes were recovered from these three tissues at weaning (**Fig. 5A**). The MLNs were the most common site of translocation, with 44% of weanlings exhibiting bacterial translocation of up to 10^5^ CFU/gram (**Fig. 5A**). In contrast, culturable bacteria were not recovered at these sites from 2-week-old PedsCom mice, suggesting that weaning-associated processes permit translocation of viable commensal microbes.

**Figure 5.**
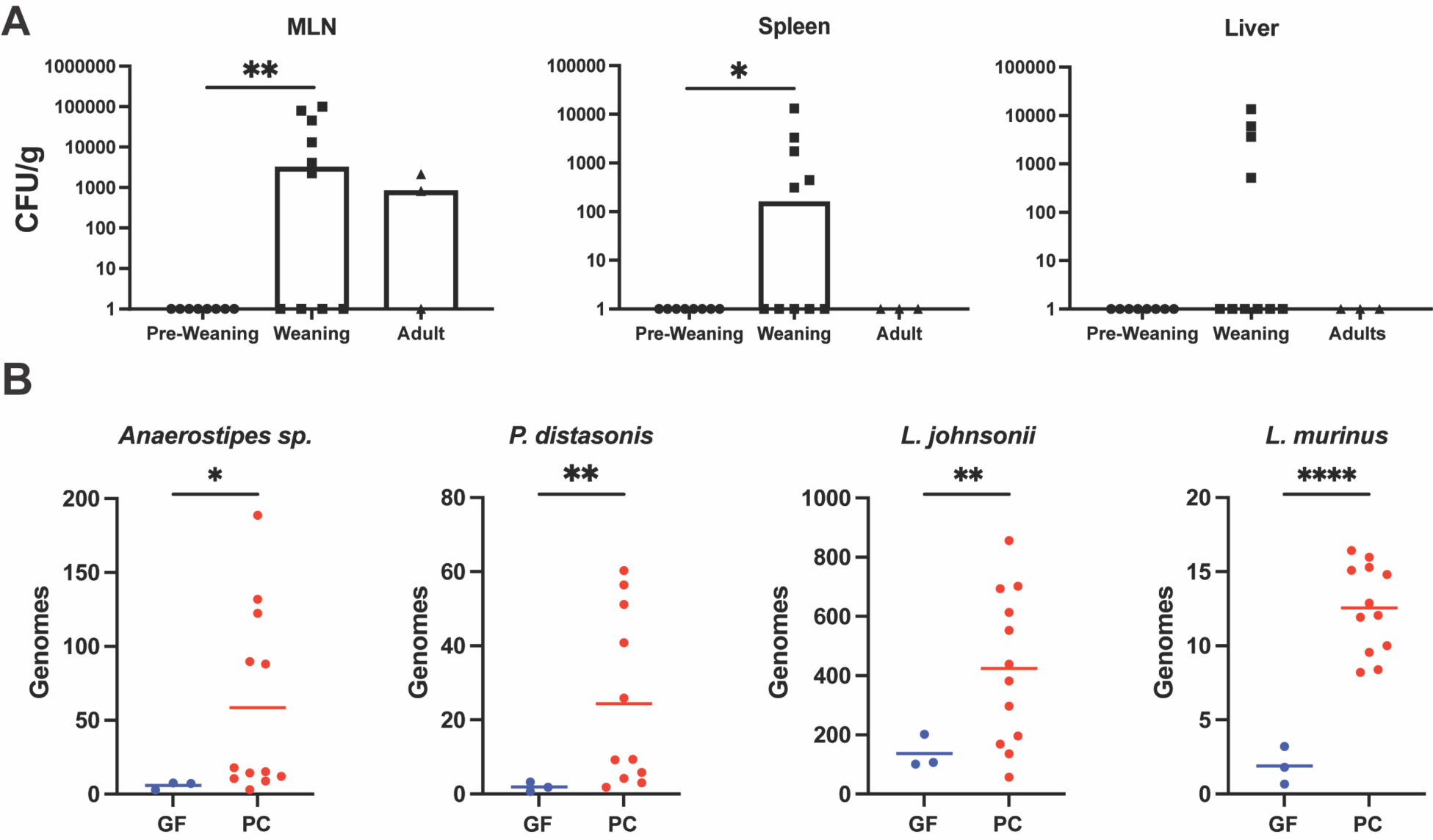
Specific PedsCom members translocate during early life. **A.** Live bacteria recovered from homogenates of mesenteric lymph nodes (MLNs), spleens, and livers of pre-weaning (2-week-old, n=8), weaning (3-week-old, n=10), and adult mice (6-week-old, n=3). Groups compared with the Kruskal-Wallis test. Representative of two independent experiments. **B.** Bacterial genomes detected in MLNs from PedsCom and germfree NOD mice by qPCR. All nine species were analyzed. The four taxa above the limit of detection and with significant differences between GF and PC samples are shown. PedsCom mice (n=11) and germfree mice (n=3). Groups compared with the Welch’s t-test. *p<0.05, **p<0.01.

To determine which PedsCom members translocate from the gut, we evaluated MLN homogenates using qPCR probes specific for the RNA polymerase beta subunit (*RpoB*) gene of each of the nine PedsCom bacteria. To ensure that intestinal samples were not contaminated during processing, we included MLNs from germfree mice as negative controls. At weaning, *P. distasonis, Anaerostipes* sp.*, L. johnsonii*, and *L. murinus* DNA was recovered from the MLNs of PedsCom NOD mice (**Fig. 5B**). In summary, of the PedsCom microbes that translocated to the MLNs, *P. distasonis,* and *Anaerostipes* sp. strongly induced pT_reg_ responses but did not induce any detectable systemic antibodies, whereas *L*. *johnsonii* induced systemic IgG1 antibodies but did not induce pT_regs_. These findings argue that bacterial translocation to systemic sites at weaning presents an opportunity for specific early-life microbes to directly influence systemic immune development. We posit that the translocation of *P. distasonis* and *Anaerostipes* sp. at weaning shapes immune ontogeny toward a regulatory state and away from autoimmunity.

## Discussion

We recently developed PedsCom, a rationally designed consortium of nine bacteria derived from the pre-weaning microbiota of diabetes-protected Eα16/NOD mice. In this study, we investigate whether this defined consortium of microbes can restrain diabetes-prone NOD mice from developing autoimmune diabetes. The two most important findings from this study are that PedsCom robustly prevents autoimmune diabetes in NOD mice and the discovery that early-life timing of colonization by PedsCom is absolutely required for this protection from T1D. In total, we identified a defined group of commensal microbes that prevents T1D, delineated a developmental window for this protection, and posit that this microbially-mediated protection may emerge at extra-intestinal sites of translocation.

PedsCom is the first tractable gnotobiotic microbial community that protects diabetes-susceptible NOD mice from disease. Previous studies demonstrated that microbes from complex undefined commensal communities can prevent the development of T1D (*31, 61, 62*). However, neither well-studied gnotobiotic communities such as altered Schaedler flora (ASF) (*63*) nor monocolonization with immunomodulatory microbes such as segmented filamentous bacteria (SFB) elicit protection from T1D (*64*). PedsCom provides a new tool to determine how commensal microbes prevent T1D. An important feature of the gnotobiotic model is that all microbial components are defined, which allows microbes to be added or removed from the community thereby defining causal relationships between specific microbes, immune populations, and disease. As proof of principle, we determined which PedsCom microbes induce regulatory T cells, and by removing these immunomodulatory microbes (*P. distasonis, Anaerostipes* sp., and *C. intestinale*), we generated a new defined community, which we called PedsCom-6. We predicted that the removal of the microbes which induce pT_regs_ would abrogate protection from T1D. Indeed, PedsCom-6 mice developed a high incidence of T1D. This microbial loss-of-function study directly demonstrates that *P. distasonis, Anaerostipes* sp., and/or *C. intestinale* are required for microbial protection from T1D.

In addition, we discovered that the timing of microbial exposure is critical for microbial protection from T1D. Seminal studies by Kasper, Blumberg, and others demonstrated that early-life colonization is required to prevent allergic disease (*19, 65*). Eberl, Hornef, and others then discovered that the early-life transition from milk to solid food, which coincides with dramatic microbiome and immune development, is crucial for maintaining long-term homeostasis (*9, 13, 55*). T1D results from an imbalance between regulatory and autoimmune processes in diabetes-susceptible individuals. Understanding what initiates and regulates this balance is critical for developing therapies to prevent T1D. Many of the early events in autoimmunity begin around weaning, including the initial priming of islet autoreactive CD4^+^ T cells, early immune cell infiltration of the islets (*66–69*), and the development of regulatory cells such as pT_regs_ (*15, 16*). We directly demonstrated that PedsCom-mediated protection from T1D is dependent on early-life commensal interactions, suggesting that there are critical microbially-driven immune interactions during this window of opportunity that restrain autoimmunity. This finding provides strong support for a causal link between specific microbes and protection from T1D, which is completely dependent on early-life exposure to these microbes. This model is consistent with human epidemiology studies of children at risk for developing T1D. For example, children who later become diabetic or develop islet autoantibodies have altered composition and functions of their gut microbiomes compared to children who do not develop T1D (*6, 70*), suggesting an influence of commensal microbes on the pathogenesis of T1D (*7, 30*). In these studies some microbes are associated with higher risk of autoimmunity while others are associated with lower risk of disease (*5, 6*). Intriguingly, *Parabacteroides distasonis* has, in some studies (*6, 53, 54, 71*), been associated with a higher risk of developing T1D, yet was associated with protection in our findings. One potential explanation is that timing of exposure to immunomodulatory microbes such as *P. distasonis* influences the trajectory of immune development toward tolerance or autoimmunity.

Introducing PedsCom microbes into diabetes-susceptible NOD mice revealed several potential pathways by which early-life commensal microbes may prevent the development of T1D. PedsCom microbes shape mucosal and systemic immune development toward tolerance by inducing regulatory T cells and potently restraining inflammation in the pancreatic islets. Specifically, we found that PedsCom colonization leads to an increase in two key IL-10-producing regulatory T-cell subsets (pT_regs_ and Tr1 cells) that restrain insulitis (*36, 37*) and T1D (*36, 38, 39*). PedsCom induces these cells more robustly than CMCom in the pancreatic lymph nodes (PLNs), which are a critical site for the initiation of islet autoimmunity and progression to T1D (*43, 44*). The pancreatic islets of PedsCom NOD mice contained far fewer IFNγ-producing effector CD4^+^ T cells than CMCom mice. Taken together, these findings argue that PedsCom microbes induce regulatory cell populations in the gut and PLNs that then restrain inflammatory T cells in the pancreatic islets. In support of this model, intestinal Tr1 cells, which can migrate to the pancreatic islets and restrain the development of T1D in an IL-10-dependent manner (*38*), were increased in PedsCom mice compared to CMCom mice. We propose that immunoregulatory cells traffic from the gut to the PLNs and pancreatic islets, where they oppose immune-mediated destruction of the insulin-producing beta cells, thereby preventing the development of T1D.

Unexpectedly, we found that *P. distasonis* and *Anaerostipes* sp. translocate to the MLN at weaning and induce pT_regs_ without inducing systemic antibody responses. In contrast, *L. johnsonii* translocates to the MLN and induces systemic anti-commensal IgG antibodies without inducing pT_regs_. These reciprocal immune responses, in which some translocating microbes induce pT_regs_ and others induce systemic antibodies, suggest that specific early-life microbes interact differently with the immune system. Gut barrier dysfunction, or a “leaky gut,” has been proposed to contribute to the pathogenesis of autoimmune diabetes based on murine (*57, 72*) and human studies of diabetic and prediabetic subjects (*73, 74*), yet the impacts of commensal translocation during early life on the development of T1D remain unclear. These unanticipated findings argue that the translocation of commensal microbes at weaning induces context-dependent immune responses, with consequences that likely depend upon the route and timing of translocation, intrinsic features of the translocating microbes, and/or the tissue environment in which immune cells and translocating microbes interact (*75*). In summary, specific early-life bacteria *(P. distasonis* and *Anaerostipes* sp.) translocate from the gut at weaning, drive tolerogenic immune responses, and contribute to early-life microbial protection from T1D.

During a short developmental stage around weaning, microbes preferentially induce Foxp3^+^RORγ^+^ pT_regs_, establishing peripheral tolerance to commensal microbes (*23, 58, 76*). This “tolerogenic window” is regulated by the complex interaction of microbial and host factors, including increasing microbiota density, changes in intestinal antigen uptake, and the development of a newly described subset of intestinal APCs (*9, 23, 77*). Live microbes at non-barrier sites typically provoke inflammation, yet here we find that the microbes that translocate at weaning in apparently healthy NOD mice induce the development of regulatory T cells in intestinal and, to a lesser extent, in systemic tissues. One possible explanation is that the host is permissive of low-level translocation of non-pathogenic microbes at weaning as a means to educate the immune system and develop tolerance toward endogenous commensal microbes. Indeed, previous studies demonstrate that microbial antigen translocation and presentation at weaning shape healthy immune development in the gut (*23*), MLNs (*78*), and thymus (*79*). In addition, a growing body of research indicates that microbial translocation influences immunity outside of the gut in the context of systemic autoimmunity (*80*) and anticancer therapies (*81–83*). Notably, two of the three PedsCom species required for protection from T1D - *P. distasonis* and *Anaerostipes sp*. - translocate to the MLNs during weaning and robustly induce pT_regs_ in intestinal and extra-intestinal tissues. These findings suggest that these microbes may promote pT_regs_ by translocating out of the gut during a tolerogenic window of immune development. Based on these findings, we propose that microbial re-localization during a pivotal developmental window may help establish tolerance and promote protection from autoimmunity. An important next step toward developing tolerogenic microbial therapeutics will be uncovering the specific microbial and host features that support the development of a diabetes-resistant immune system.

In summary, we designed a simple consortium of nine pre-weaning microbes that prevents T1D during a crucial early-life window of opportunity. These findings provide a framework for understanding how factors that shape the intestinal microbiome early in life, such as mode of birth, diet, and antibiotic treatments, may alter the risk of developing autoimmunity later in life (*84*). The time-dependent immune effects of PedsCom colonization argue that we should carefully control for timing when designing and testing microbial therapeutics to prevent immune-mediated diseases. Our results provide compelling evidence that early-life microbial interventions targeting a critical window of immune ontogeny have the potential to prevent T1D in susceptible populations.

## Methods and materials

### Mice

Germfree NOD mice were generated by Caesarian-section delivery and cross-fostered to germfree Swiss-Webster mice. Gnotobiotic mice were bred and housed in the Hill Pavilion vivarium at the University of Pennsylvania. Mice were fed the Autoclaved LabDiet 5021 (Cat# 0006540) *ad libitum* and housed on autoclaved Beta-chip hardwood bedding (Nepco, NY, USA). To generate the PedsCom NOD gnotobiotic line, germfree 6 to 8-week-old NOD mice were gavaged with 10^9^ CFU of each of the 9 PedsCom species in 100 mL of reduced PBS (rPBS). To generate the CMCom gnotobiotic line, whole cecal contents of a 6-week-old female SPF Eα16/NOD mouse were collected anaerobically in 5mL rPBS, the contents settled for 10 mins, and then 100 mL of the supernatant was used as donor microbiota material for oral gavage (*13*). Gavaged mice were bred in separate PedsCom and CMCom gnotobiotic isolators, then F1 and later generations were used experimentally, to ensure natural vertical transmission of microbiota. Sterility checks were regularly performed on germfree isolators each month. Freshly collected pellets were cultured on brain heart infusion (BHI) (Oxoid, UK), NB1, and Sabouraud media for 65–70 hours at 37°C under aerobic and anaerobic conditions with positive and negative control samples. Isolator sterility was confirmed externally every 3-4 months by Charles River Laboratories (NJ, USA). PedsCom gnotobiotic line was confirmed by 16sRNA gene metagenomic sequencing every 3-4 months. Unless otherwise indicated, PedsCom and CMCom colonized NOD mice acquired their microbiota via vertical transmission from their dams at birth.

For monocolonization experiments, NOD mice at 3 weeks of age were orally gavaged with 50 mL of rPBS, 10^8^ CFU of individual PedsCom species in 50 mL of rPBS, or 50 mL of cecal contents of 8-week-old female PedsCom NOD mouse that was collected from a 5 mL slurry in rPBS. Gavages were performed using a sterile syringe and a straight 1-inch-long 22-gauge gavage needle (Cadence Science, #7901). For post-weaning colonization experiments, 6-week-old germfree NOD females were transferred to the PedsCom gnotobiotic isolator and cohoused with female PedsCom mice. All mouse experiments were approved by the Institutional Animal Care and Use Committee (IACUC) at the University of Pennsylvania.

### Bacteria

Individual PedsCom species were isolated and cultured from 14-day-old Eα16/NOD mice, as previously described (*13*). *Clostridium intestinale, Anaerostipes* sp., and *Parabacteroides distasonis* were grown anaerobically at 37°C in brain heart infusion (BHI) media (Oxoid, UK) supplemented with hemin (5 mg/L) and vitamin K (0.5 mg/L) (BD, Franklin Lakes, NJ). *Lactobacillus johnsonii* and *Ligilactobacillus murinus* were grown anaerobically at 37°C in de Man Rogosa Sharpe (MRS) media (Sigma, MO, USA). *Staphylococcus sciuri, Kosakonia cowanii, Enterococcus faecalis,* and *Staphylococcus xylosus* were grown in ambient air at 37°C in BHI media. Liquid cultures were centrifuged at 4700 x g at 4°C and resuspended in sterile reduced, phosphate-buffered saline containing 0.1% L-cysteine (rPBS). The optical density (OD) of cultures was determined using a spectrophotometer and the bacteria concentration was adjusted to an O.D. of 0.1 for microbial flow cytometry experiments and 10^8^ colony-forming units (CFU) for experiments requiring monocolonization.

### Diabetes and insulitis

To assess diabetes in gnotobiotic mice, mice were screened for glucosuria starting at 10 weeks of age. Urine was collected and analyzed for glucosuria by dipstick (Diastix; Ames). After two consecutive positive tests for glucosuria, diabetes was confirmed by blood glucose greater than 250 mg/dL. Mice were monitored until 30 weeks of age. To assess insulitis, pancreata were harvested at 10 weeks of age and fixed in 10% formalin. Then pancreata were embedded in paraffin, sectioned, and stained with hematoxylin and eosin. Insulitis was evaluated by a blinded researcher using light microscopy. Islets were scored 0 for no signs of insulitis, 1 for peri-insulitis, and 2 for insulitis. Each mouse was given an average composite score of insulitis (range 0-2).

### Microbial flow cytometry (mFLOW)

Cultured bacteria were centrifuged at 4700 x g for 10 minutes, resuspended in 1 mL PBS, 1% BSA (PBS-BSA), and diluted to an OD600 of 0.1. Bacteria were washed with PBS-BSA, resuspended in blocking buffer (PBS-BSA, 20% normal rat serum), and incubated for 30 minutes at 4°C. The bacteria were then incubated with serum (heat inactivated at 56°C for 30 min, centrifuged at 16000 x g for 5 min at 4°C) diluted 1:25 in bacterial staining buffer (BSB) (PBS, 1% BSA plus sodium azide) and incubated at 4°C for 1 hr. The bound bacteria were then washed with BSB and incubated with anti-mouse antibodies PE-Cy7-IgG1, clone RMG1-1; BV711-IgG2b, clone R12-3; BV421-IgG3, clone R40-82) diluted 1:25 in BSB for 30 minutes at 4°C, washed and incubated in SytoBC nucleic acid stain (Invitrogen #S34855) diluted 1:500 in Tris-buffered saline (TBS) for 15 minutes at room temperature. Flow cytometry analysis on stained bacteria populations with SytoBC was used to define living bacteria. RAG2 KO sera was used to define gates for Ig-coated bacteria and control for nonspecific binding of secondary antibodies. Notably, *L. murinus* demonstrated nonspecific binding of secondary antibodies and was excluded from further analysis. PedsCom samples were compared to sera from germfree mice to determine if the antibody response was elicited from colonization rather than natural antibodies. Microbial flow cytometry analysis was completed on the LSR Fortessa and data analysis was performed using FlowJo v10 software (BD).

### Reverse transcription quantitative PCR

Fecal pellets or intestinal contents were collected in sterile microcentrifuge tubes and stored at −80°C. Bacterial DNA was isolated from samples using the DNAeasy PowerSoil kit (Qiagen, Germany) using a QIACube (Qiagen, #9001292). PedsCom species abundance was determined by qPCR, using TaqMan Master Mix (Thermo-Fisher Scientific, Waltham, MA) and primers for the DNA-directed RNA polymerase subunit beta (*rpoB*) gene for each specific PedsCom species. Primer and probe sets were generated using the PrimerQuest Tool (Integrated DNA Technologies, IA, USA) and compared with the Multiple Primer Analyzer tool (Thermo-Fisher Scientific, Waltham, MA) (*13*). A standard curve using serial dilution of cultured bacteria was used to estimate the genomes of each PedsCom species in a sample and determine the limit of detection of the assay for each PedsCom member.

### 16S rRNA gene metagenomic sequencing

The PennCHOP microbiome program sequencing core sequenced the V4 variable region of the 16S rRNA gene using the Illumina MiSeq platform as previously described (*85*). QIIME2 (ver. 2018.2) was used to analyze the resulting sequencing libraries and deblur was used to cluster and de-noise the results into amplicon sequence variants (*86*). The Greengenes 99% rRNA gene reference database (ver. 13.8) was used to classify bacterial taxa. Taxa bar plots were generated with the R package phyloseq (*87*).

### Isolation of immune cells from tissues

Mouse spleen, MLN, PLN, cecum, and large intestine tissues were isolated from euthanized mice and placed in cell culture media (CCM: RPMI, 100 units/mL penicillin, 100 μg/mL streptomycin, 10% fetal bovine serum (FBS)). The lymph nodes and spleen were dissociated using the back of the plunger of a 1 mL syringe through a 70 μM filter in 1 mL CCM in a dish. The dish and filter were rinsed with CCM, and the suspension was spun down at 480 x g for 10 minutes in 15 mL conical tubes. The supernatant was discarded, and MLNs and PLNs pellets were resuspended in 200 μL CCM and plated in a 96-well round-bottom plate for staining. Splenocytes were resuspended in 1 mL ACK lysis buffer for 1 minute and then quenched with 10 mL CCM and centrifuged as before. The pellet was then resuspended in 1 mL staining media (RPMI, 4% FBS) and 100 μL of each sample was plated for staining. The intestines were cleared of debris by gently flushing with cold PBS using a gavage needle or by cutting open the cecum and shaking in CCM. The cecal patch was carefully excised. The colon was rolled on a paper towel and a scalpel was used to remove mesenchyme. All tissues were then cut open and shaken to remove the remaining intestinal debris. The intestinal tissues were then incubated in intestinal epithelial layer (IEL) digestion media (30 mL RPMI, 0.062% dithiothreitol, 1 μM EDTA, 1.3% FBS) and incubated at 37°C for 15 minutes while stirring at 300 rpm with a magnetic bar to remove IEL cells. The tissues were washed with PBS, placed in RPMI, and diced into small sections (∼1 mm in size) using dissection scissors. The tissues were then digested for 40 minutes in 25 mL digestion media (25 mL RPMI, 12.5 mg dispase (Gibco), 37.5 mg collagenase type II (Gibco), 1 mL PBS) at 37°C with stirring at 300 rpm to generate a single cell suspension. The suspension was filtered through 100 μM filters and quenched with 25 mL RPMI. The solutions were centrifuged for 10 minutes at 310 x g at 4°C and resuspended in FACS buffer (DPBS, 0.1% BSA, 2 mM EDTA).

The pancreas was perfused with 0.5 mL of C-solution (HBSS, 0.35g/L NaHCO_3_, 2.5 % bovine serum albumin (BSA), collagenase P 2 mg/mL (Roche, Mannheim, Germany, #11213857001), then carefully removed. The pancreas was incubated in 10 mL C-solution in a water bath at 37° C for 12 minutes. The pancreas tissue was hand-shaken vigorously for 30 seconds to break up the loose tissue. The pancreas tissue was washed three times with 15 mL cold G-solution (HBSS, 0.35g/L NaHCO_3_, 2.5 % BSA) to remove the collagenase by spinning down at 4°C at 220 x g for 5 minutes. The tissue homogenate in 50 mL cold G-solution was then filtered through a size 40 sieve (425 um diameter wire mesh, Bellco Glass Cat # 1985-00040) to remove the remaining undigested tissue, fat, and lymph. The tissue homogenate was then centrifuged at 4°C at 220 x g for 5 minutes and resuspended in 1.1g/mL islet suspension by mixing 7.5 mL Histopaque 1.119 (Sigma) with 3.5 mL Histopaque 1.077 (Sigma). 7 ml of Histopaque 1.119 was added under the islet phase using a needle. 12 ml Histopaque 1.077 was overlayed, and then 12 mL of G solution was overlayed on the Histopaque gradient. The gradient solution was then centrifuged for 25 minutes at 1750 x g with very slow acceleration and no braking (1/0) at 4°C. The islet layer was collected from each of the interfaces by aspirating 10 mL at each interface. The islets were then washed with 50mL G-solution and centrifuged at 220 x g for 5 minutes to remove Histopaque. The islets were then incubated with 5 mL of 0.1 mM EDTA RPMI solution for 5 minutes. The islets were then washed again and resuspended in 5 mL of pre-warmed dispase solution (1mg/mL in RPMI). The islets were incubated for 15 minutes at 37°C to dissociate islets into a single-cell suspension. The samples were then centrifuged at 4°C at 220 x g for 5 minutes and resuspended in FACS buffer in preparation for staining.

### Flow cytometry of immune cells

Staining for surface markers of immune cells was carried out in the dark at 4°C for 15 – 30 minutes using the following antibody panel: BV510-CD45 (clone 30F11; Biolegend, CA, USA), PE-Cy7-TCRβ (clone H57-597; Biolegend), APC-Cy7-CD19 (clone 6D5; Biolegend), AF700-CD8 (clone 53-6.7; Biolegend), FITC-CD4 (clone Rm4-5; Biolegend). Cells were washed twice with staining media and fixed overnight at 4°C using the eBioscience Fixation kit (cat# 005223-56, 00-5123-43). Cells were washed twice in permeabilization buffer (Invitrogen cat# 00-8333-56), then centrifuged at 1340 x g and stained for intracellular proteins for 50 minutes at room temperature using the following panel: APC-Foxp3 (clone FJK-16s; eBioscience, MA, USA), PE-RORγ (clone B2D; eBioscience), Pacific Blue-Helios (clone 22F6; Biolegend). Cells were washed twice with permeabilization buffer, resuspended in 200 μL of PBS, and filtered through 50 μM nylon mesh (Genesee Scientific, cat# 57-106) into FACS tubes. To detect intracellular cytokines, cells were re-suspended in 200 μL of cell stimulation cocktail with protein transport inhibitors (Thermo-Fisher Scientific, Cat: 00-4975-03) in CCM for 2 hours at 37°C and 5% CO_2_. Cells were then washed, resuspended, surface stained, and permeabilized as described above. The samples were then intracellularly stained with IL-10-BV421 (clone JES5-16E3; Biolegend) and IFNγ-AF488 (clone XMG1.2; Biolegend) as described above. The cells were washed and then resuspended in 500 µl FACS buffer for analysis. Stained samples were analyzed on the LSRFortessa (BD) or Aurora CyTek and data analysis was performed using FlowJo v10 software (BD).

### Microbial translocation

MLNs, spleen, and a segment of liver were sterilely harvested and placed into autoclaved pre-weighed 1.5-mL Eppendorf tubes. The tissue weights were then recorded. Tissue samples were then processed in an anaerobic chamber (90% N_2,_ 5% CO_2,_ and 5% H_2_). Samples were homogenized in sterile rPBS. Portions of the tissue homogenates were then inoculated onto Yeast Casitone Fatty Acids with Carbohydrate (YCFAC) plates, then incubated anaerobically at 37°C for 48 hours. After incubation, colony-forming units (CFUs) were counted and normalized to the weight of tissue (CFU per gram). A pseudo-count of +1 was used for samples with no growth to visualize the sample on a log axis. A portion of the sample tissue homogenate was used to identify and quantify individual PedsCom species by quantitative real-time PCR. Additionally, samples were compared to tissue samples from germfree control mice to further validate the assay.

### Bioinformatics and statistics

Statistical analysis was completed using Prism 10. For diabetes studies, a Log-rank test was used to compare groups. For parametric analysis of more than two groups, the ANOVA test on mean values was used with the Holm-Sidak multiple comparisons test for comparisons of all means within a test group to correct for multiple comparisons. For non-parametric analysis of more than two groups, the Kruskal-Wallis test on median values was used with post-hoc analysis consisting of the uncorrected Dunn’s test for multiple comparisons of two medians within the Kruskal-Wallis test. For the non-parametric analysis of two groups, the Mann-Whitney-Wilcoxon test on median values was used. For parametric analysis with an unequal variance of two groups, the Welch’s t-test on mean values was used. For multivariate analysis, the two-way-ANOVA test with posthoc analysis consisting of the Fisher’s LSD test for direct comparisons of two means within the two-way-ANOVA test was used.

## List of Supplementary Materials

Figure S1. Microbiome of the Complex Mature Community CMCom.

Figure S2. PedsCom induces IL-10-producing T cells.

Figure S3. Similar levels of insulitis in PedsCom and CMCom NOD mice.

Figure S4. Specific PedsCom species induce systemic antibodies.

Figure S5. Anaerostipes sp. and P. distasonis induce pTregs and do not elicit IgG1.

## Acknowledgments

The authors thank Drs. Nissan Yissachar, Neil Surana, Ike Eisenlohr, and Will Bailis for their helpful input. The authors also thank Dmitri Kobuley and Michelle Albright of the PennCHOP Microbiome Program gnotobiotic mouse facility. Some images were generated with Biorender.

## Author contributions

Conceptualization, J.G. and M.A.S.; Methodology, J.G. and M.A.S.; Investigation, J.G., J.D., J.N.F., S.M., T.D., L.G. and M.A.S. Writing – Original Draft, J.G. and M.A.S.; Writing – Review and Editing, J.G., J.D., J.N.F., S.M., T.D., L.G., P.J.P., L.C.E and M.A.S.; Funding Acquisition, J.G. and M.A.S.; Resources, M.A.S.; Supervision, M.A.S. and L.C.E.

