## Supplemental Figures for "Early-life exposures to specific commensal microbes prevent type 1 diabetes"

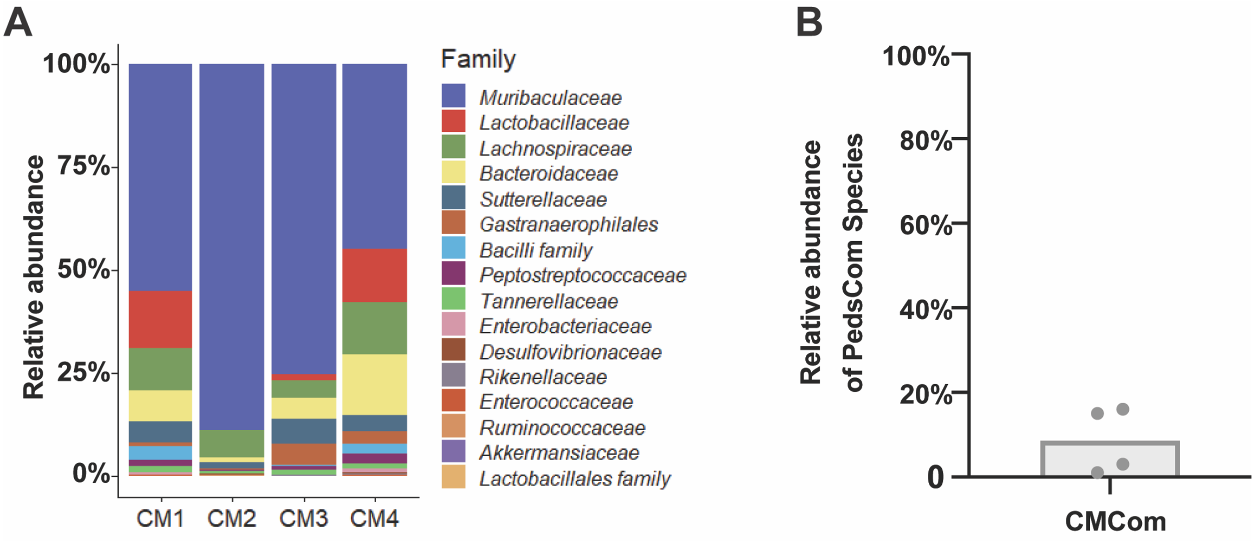


**Figure S1. Microbiome of the Complex Mature Community (CMCom). A.** Representative family level taxonomy of fecal microbiota of NOD mice colonized with the CMCom community. CM1, CM2, CM3, CM4 denote individual mice. **B.** Mean read coverage of PedsCom taxa in fecal microbiome of 6-week-old adult CMCom NOD mice (n=4).

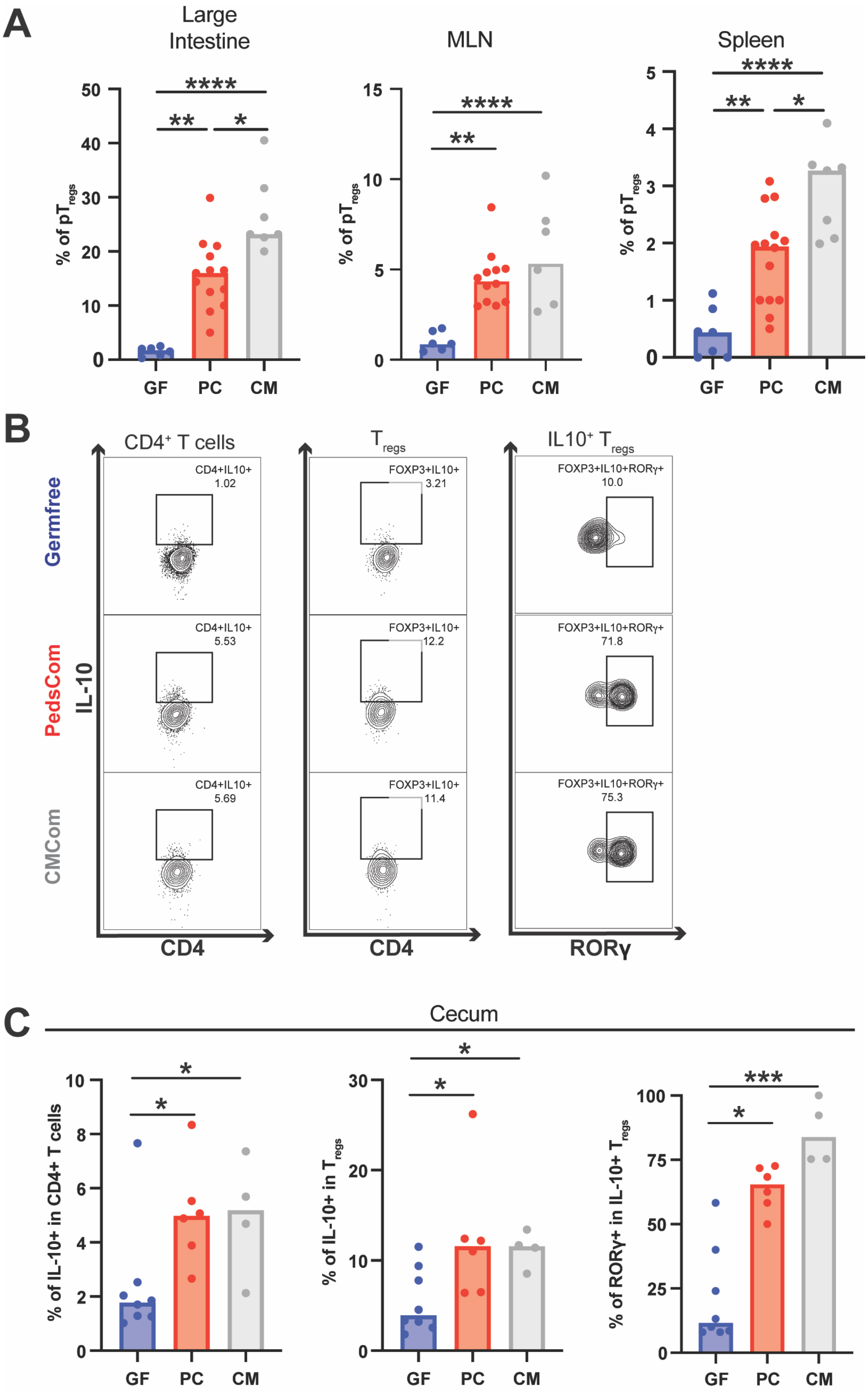

A
Large Intestine
MLN
Spleen
% of pTregs
****
**
*
GF
PC
CM
B
CD4+ T cells
Tregs
IL10+ Tregs
Germfree
PedsCom
CMCom
IL-10
CD4+IL10+
1.02
5.53
5.69
FOXP3+IL10+
3.21
12.2
11.4
FOXP3+IL10+RORγ+
10.0
71.8
75.3
CD4
RORγ
C
Cecum
% of IL-10+ in CD4+ T cells
% of IL-10+ in Tregs
% of RORγ+ in IL-10+ Tregs
***

**Figure S2. PedsCom induces IL-10-producing T cells. A.** Percentages of $pT_{regs}$ ($CD4^+FOXP3^+ROR\gamma^+Helios^-$) of total $FOXP3^+$ $T_{regs}$ in the large intestines, mesenteric lymph nodes (MLNs), and spleens of germfree, PedsCom, and CMCom NOD mice. n≥4 samples per tissue per gnotobiotic community, ages 5-11 weeks old. Groups compared with the Kruskal-Wallis test.**B.** Representative flow cytometry of IL-10 producing $CD4^+$ T cells, $CD4^+FOXP3^+$ $T_{regs}$ and $ROR\gamma^+$IL-10 producing $T_{regs}$. **C.** Percentages of IL-10 producing $CD4^+$ T cells in the ceca of germfree, PedsCom and CMCom mice. Percentages of IL-10 producing $T_{regs}$ in the ceca of germfree, PedsCom and CMCom mice. Percentages of $ROR\gamma^+$ $T_{regs}$ in IL-10 producing $T_{regs}$ in the ceca of germfree, PedsCom and CMCom mice. n ≥3 tissues per gnotobiotic community, ages 10-12 weeks. Groups compared by the Kruskal-Wallis test, *p<0.05, **p<0.01, ***p<0.001, ****p<0.0001. GF=germfree, PC=PedsCom, CM=CMCom.

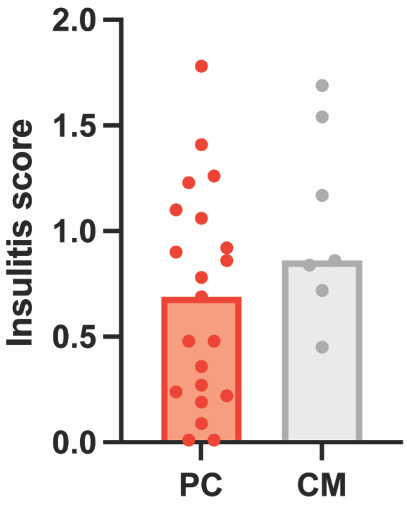


**Figure S3. Similar levels of insulitis in PedsCom and CMCom NOD mice. A.** Composite insulitis score from NOD female mice at 10-12 weeks of age. n>7 tissues per gnotobiotic community. Groups compared with the Mann-Whitney-Wilcoxon test.

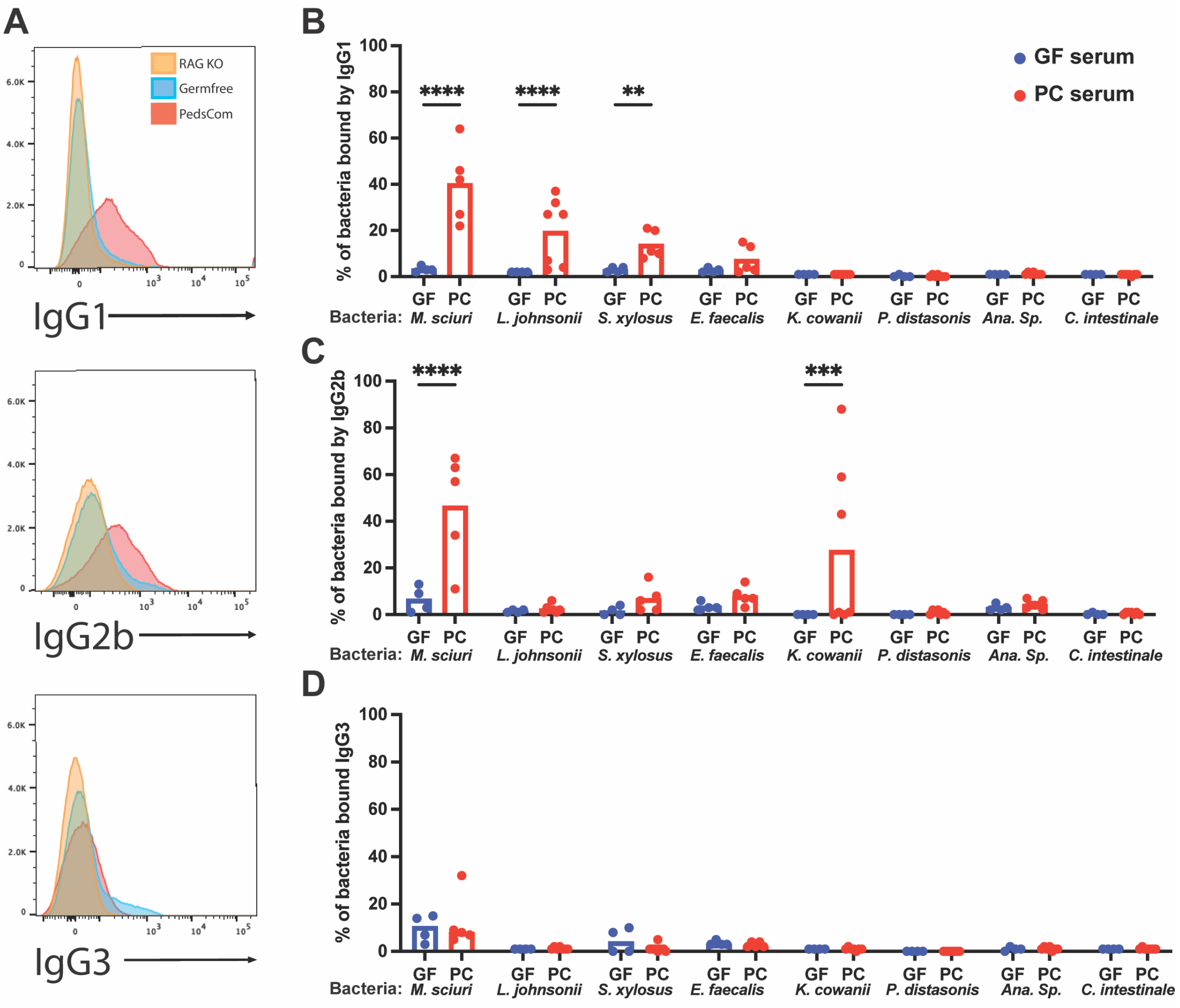


**Figure S4. Specific PedsCom species induce systemic antibodies. A.** Sera from germfree, PedsCom, and RAG2 deficient mice were incubated with pure cultures of individual species from PedsCom**.** Histogram of microbial flow cytometry of systemic IgG1, IgG2b, IgG3 binding to *M. sciuri*. **B-D.** Frequency of each PedsCom species bound by IgG1 (B), IgG2b (C), or IgG3 (D) from serum of germfree or PedsCom mice. *L. murinus* was excluded due to high levels of nonspecific binding of secondary antibody. Groups compared by the two-way ANOVA test**,** n≥5 PedsCom mice and n=4 germfree mice, **p<0.01, ****p<0.0001. GF=germfree, PC=PedsCom.

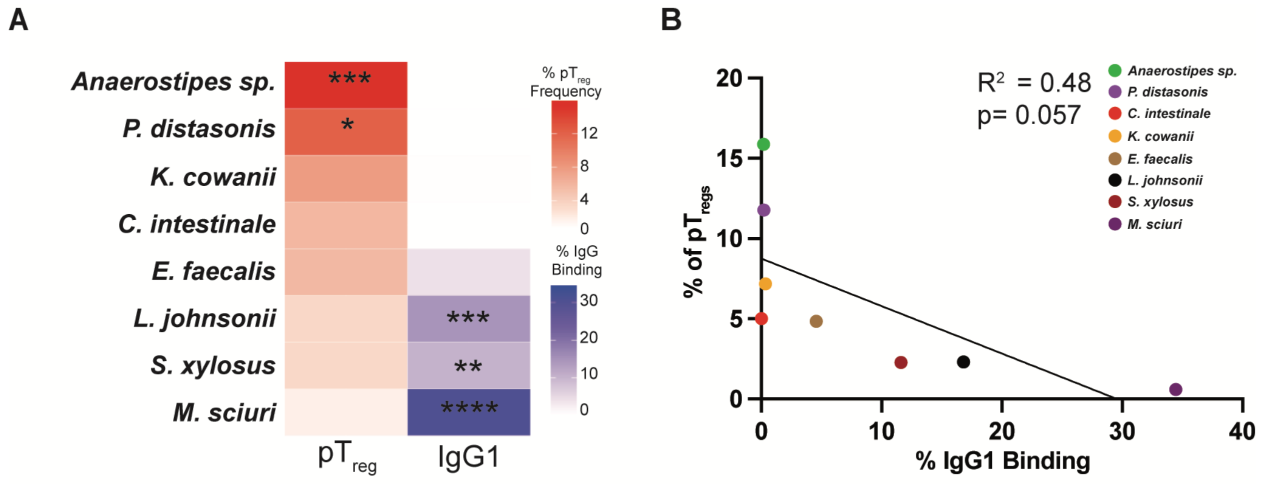


**Figure S5. Anaerostipes sp. and P. distasonis induce pTregs but do not elicit IgG1. A.** Heat map depicting the mean $pT_{regs}$ ($CD4^+FOXP3^+ROR\gamma^+Helios^-$) percentages of total $FOXP3^+$ $T_{regs}$ in the large intestines of individual PedsCom monocolonized mice and the mean binding of systemic antibodies to individual PedsCom species. Groups compared by the ANOVA test, n≥3 mice per PedsCom microbe **C.** Linear regression of $pT_{reg}$ frequency and IgG1 binding to individual PedsCom species. *p<0.05, **p<0.01 ***p<0.001, ****p<0.0001

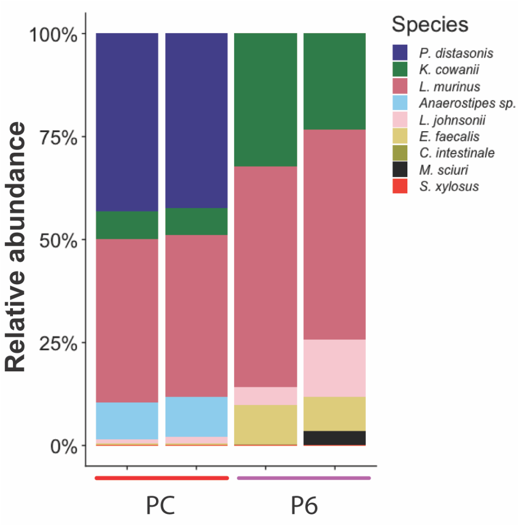


**Figure S6. Microbiome of PedsCom-6 (P6) community.** Relative abundance of PedsCom members in feces of P6 and PC (PedsCom) NOD mice.
